# A Full Window Data Independent Acquisition Method for Deeper Top-down Proteomics

**DOI:** 10.1101/2024.11.08.622616

**Authors:** Chen Sun, Wenjing Zhang, Mowei Zhou, Martin Myu, Wei Xu

## Abstract

Top-down proteomics (TDP) is emerging as a vital tool for the comprehensive characterization of proteoforms. However, as its core technology, top-down mass spectrometry (TDMS) still faces significant analytical challenges. While data-independent acquisition (DIA) has revolutionized bottom-up proteomics and metabolomics, they are rarely employed in TDP. The unique feature of protein ions in an electrospray mass spectrum, as well as the data complexity require the development of new DIA strategies. This study introduces a machine learning assisted Full Window DIA (FW-DIA) method that eliminates precursor ion isolation, making it compatible with a wide range of commercial mass spectrometers. Moreover, FW-DIA leverages all precursor protein ions to generate high-quality tandem mass spectra, enhancing signal intensities by ∼50-fold and protein sequence coverage by threefold in a modular protein analysis. The method was successfully applied to the analysis of a five-protein mixture under native conditions and *Escherichia coli* ribosomal proteoform characterization.

## Introduction

Proteoforms represent the diverse array of protein species generated from a specific gene, that play a crucial role in biological evolution and regulation of life activities^[1]^. By analyzing intact proteins without digestion, top-down proteomics (TDP) offers a comprehensive perspective of the proteome and has been extensively employed for proteoform characterizations over the past few decades^[1–3]^. Recent advancements in front-end protein preparation and separation^[4–6]^, mass spectrometry (MS) instrumentation^[7–9]^, and bioinformatics^[11–12]^ have significantly enhanced the capability of TDP, thereby expanding its applications in the realm of biology,^[13]^ biomedical and pharmaceutical researches^[14–16]^.

Currently, data-dependent acquisition (DDA)^[17]^ method is the predominant data acquisition technique used in TDP^[18–21]^. However, traditional DDA methods face challenges due to the stochastic nature of precursor ion selection, leading to difficulties in detecting low-abundance ions^[22]^. Moreover, protein ions generated by electrospray ionization (ESI) typically possess multi-charge states. The precursor ion isolation only utilizes a small portion of ions generated by a given protein molecule, leading to diminished fragment ion intensities. This scenario contrasts with that of small ions in metabolomics, where a specific small molecule usually has only a single charge state (e.g., 1+). Consequently, the DDA method in TDP may exhibit low reproducibility and is prone to lose information. Such limitations can compromise the accuracy and comprehensiveness of proteomic analysis, potentially resulting in an incomplete or biased depiction of the biological system under study.

To overcome these challenges, the adoption of data-independent acquisition (DIA)^[23]^ methods becomes a natural progression, which have revolutionized the field of bottom-up proteomics (BUP). DIA technologies have become a favored approach in BUP given their high throughput, unbiased MS analysis, exceptional consistency, and ability to accommodate low-abundance proteins^[24]^. Several DIA methods have been developed specifically for BUP^[25]^. These can be classified into two main categories: spectrum-centric (such as DIA-Umpire^[26]^, Group-DIA^[27]^) and peptide-centric methods (such as DreamDIA^[28]^, MaxDIA^[29]^, DIA-NN^[30]^, OpenSWATH^[31]^, Spectronaut^[32]^ and Skyline^[33]^)^[34]^. While these tools have been successful in BUP, adapting these tools for TDP is not straightforward. This is because proteoform identification in TDP differs fundamentally from the identification of proteolytic peptides in BUP. First, an intact protein in an ESI mass spectrum usually shows a broad mass range distribution with multiply charged ions. The traditional DIA isolation windows used in BUP may not be wide enough to capture all charge state ions, resulting in information loss and diminished fragment signals. Furthermore, the introduction of DIA in TDP will significantly increase the complexity of data analysis. This complexity arises not only from the unique characteristics of TDP data, such as high charge states, large molecular weights and spectral overlap, but also from the complex fragment ion spectra and the loss of precursor-fragment ion relationships introduced by DIA approaches^[35,36]^. Existing TDP software tools, such as ProSightPD, TDPortal (http://nrtdp.northwestern.edu/resource-software/), Informed-Proteomics^[37]^, ProSight Lite^[38]^, TopPIC^[39]^, and pTop^[40]^ are primarily designed for processing DDA data.

Developing DIA methods specifically designed for TDP is crucial to unlock its full potential and overcome the limitations of traditional DDA methods. In this study, we introduce a novel DIA technique called full window DIA (FW-DIA). In contrast to TopDIA^[41]^, FW-DIA involves fragmenting all protein ions eluted from the liquid chromatography (LC) system without performing ion isolation. By doing so, we can maximize the use of precursor protein ions, enhancing both fragment ion intensities and the variety of fragment ion species. To establish precursor-fragment ion relationship and minimize interference, a machine learning assisted data processing scheme was developed in FW-DIA, which includes feature detection and ion pairing strategy to correlate precursor protein molecules with their respective fragments. Additionally, an isotope ratio-based verification module was implemented to perform quality control and validate results. This FW-DIA was characterized using model proteins, and applied for the analysis of *E. coli* ribosomal proteins. FW-DIA consistently outperformed DDA in terms of protein identification, sequence coverage, and localization of post-translational modifications (PTMs). Besides deepening proteoform identification, FW-DIA enables the application of DIA-TDP on a wide range of commercial MS instruments, which would facilitate the fast evolution and broad application of TDP.

## Results

### Overview of the FW-DIA procedure

In the FW-DIA experiment, proteins eluted from LC are analyzed by the mass spectrometer alternating between two functions. The first function (the survey scan) acquires low-energy (6 eV) precursor ion mass spectra. In the second function (MS/MS scan), all ions in the mass range would undergo high energy collision dissociation (HCD) in the collision cell, and elevated-energy fragment ion mass spectra are then collected (Figure 1a). The data processing pipeline for FW-DIA consists of three key components: data preprocessing, feature detection and ion pairing. As shown in Figure 1b, the raw data undergo initial preprocessing using a convolutional neural network (CNN)-based model to distinguish genuine protein signals from noise (Supplementary Fig. 1-2). This preprocessing step is critical for reducing noise and ensuring accurate downstream analysis in TDP. The CNN model was trained using ESI mass spectra data collected on the same mass spectrometer. Spectral deconvolution was carried out with FLASHDeconv^[42]^, and the resulting deconvoluted spectra were manually reviewed to accurately label protein signals and noise, ensuring high-quality training data. The CNN model was then trained as a binary classification task to classify protein peaks from noise. The preprocessing framework utilizes two specialized models, one for MS1 and one for MS2 data, to handle the distinct characteristics of each dataset. As demonstrated in Supplementary Fig. 3, the noise density level could be reduced from 10^5^ to 10^4^ in a mass spectrum. This denoising process significantly lowered the false positive rate in the subsequent deconvolution step, improving both accuracy and reliability. Furthermore, the deep learning-based denoising method facilitated the deconvolution of a broader and more comprehensive range of masses. This resulted in a substantial increase in the number of deconvoluted masses by up to 28.4%, and a nearly doubled mass coverage range in the deconvolution step (Supplementary Fig. 4).

**Figure 1.**
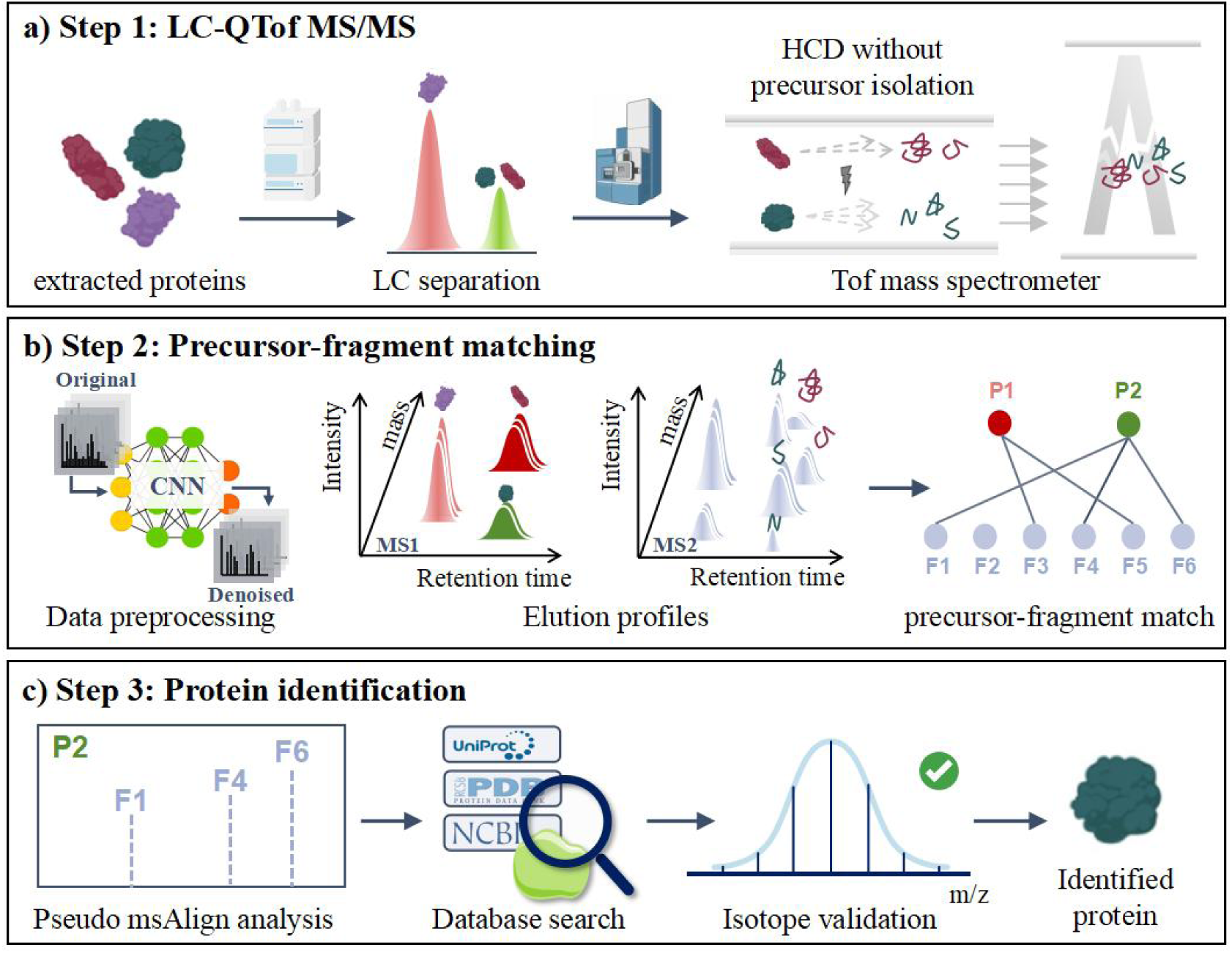
Schematic diagram of experimental flowchart. a) Step 1: Schematic diagram of the LC-QTof experiment setup; b) Step 2: Precursor-fragment ion matching; c) Step 3: Proteoform identification and verification.

After denoising, the elution profiles of precursors and fragments are then built by the feature detection module; while the ion pairing module matches fragments with their precursors based on co-elution feature behavior to establish precursor-fragment relationships. In typical TDP, spectra are first deconvoluted to simplify the data by merging signals across isotopes and charge states. During deconvolution, the feature detection module identifies and extracts a complete set of features, enabling the generation of elution profiles for both proteoforms and their corresponding fragments. The accuracy of deconvolution results is critical for reliable proteoform identification^[43–45]^. To obtain comprehensive and accurate feature detection, this procedure also integrates the deconvolution results from different deconvolution pipelines. Three distinct deconvolution tools were chosen: Unidec (6.0.4 version)^[46]^, FLASHDeconv (3.0 version)^[42]^ and TopFD^[47]^. Since different deconvolution strategies were applied, these three methods are complementary to each other. By analyzing the output of these tools, molecular weight, continuous m/z value, and retention-time span were used to determine whether the results from each tool pointed to the same entities.

Following feature detection, which identifies all possible precursor and fragment ion signals, ion pairing is used to match precursor-fragment pairs. To assess the likelihood that a detected fragment signal originates from a specific precursor proteoform ion, the algorithm first selects fragment candidates that co-elute with the precursor within a tolerated scan time range. Next, the algorithm calculates the Pearson correlation coefficient (PCC) of the LC elution profiles between the detected precursor features and the fragment ion candidates. Sets of fragment peaks (with PCC > 0.8) are grouped with precursor features and stored as precursor-fragment groups. Finally, the algorithm generates a pseudo-msAlign file (.msalign) that records the relationships between precursors and fragments. For each precursor, the msAlign file includes all possible fragments of the intact protein, providing a comprehensive representation of proteoform composition.

In step 3 (Figure 1c), we employed TopPIC^[39]^ as the search engine for protein identification. TopPIC accepts deconvolved spectra in the msAlign file format, a simplified text format derived from Mascot Generic Format. This format typically records precursor and fragment ion information extracted from DDA experiments. Due to the similarities between DDA and DIA pseudo-msAlign files, subsequent analysis of database search results can be performed comparably. The pseudo-msAlign file, along with a protein database file in FASTA format, serves as the input for TopPIC, which then identifies protein proteoforms.

After obtaining results from TopPIC, a verification process was implemented to validate the identified protein fragments and assess data quality. This process included the alignment of observed fragment isotope peaks with their corresponding theoretical counterparts. The degree of similarity between experimental and theoretical isotopic envelopes was quantified using PCC, and a threshold of 0.8 was selected to serve as an essential quality control metric for accurate fragment matching. Theoretical isotopic envelopes for the fragment ions were calculated using the R package enviPat^[48]^, which utilized the chemical formula and charge state of each fragment ion as input parameters. Experimental isotopic envelopes were directly extracted from output of the feature detection stage. To ensure that intensity and m/z position information were comparable, both experimental and theoretical isotopic envelopes were normalized prior to calculate isotope similarity (see Supplementary Fig. 5).

### Analysis of a model protein

Performances of the FW-DIA method was first evaluated and compared with the conventional TD-DDA approach using a model protein cytochrome C (Cyt C). The results of DIA and DDA were compared after optimizing the experimental conditions separately (see Supplementary Fig. 6). In a typical ESI mass spectrum, a protein would possess multiple charge states, as well as multiple isotope peaks within each charge state. In DDA experiments, precursor ions need to be first isolated, then undergo fragmentation. For protein samples, typically protein ions of a selected charge state (within the isolation window) were isolated and then analyzed in a single MS/MS scan; while ions of other charge states were discarded. This TD-DDA operation would cause two problems: 1. Only a subset of protein ions was utilized, lowering intensities of fragment ions; 2. Protein ions at different charge states would possess different conformations, with ions at higher charge states generally adopting more unfolded structures^[50]^. The fragmentation patterns of protein ions are influenced by their conformations, meaning that ions at different charge states will produce different fragment ions and generate complementary tandem mass spectra. Therefore, each tandem mass spectrum contains a limited number of fragment ions, which reduces the protein sequence coverage rate. On the other hand, the FW-DIA fragments all protein ions in a single tandem MS experiment, resulting in an enriched set of fragment ions with higher intensities.

As shown in Figure 2a, the tandem mass spectrum of Cyt C collected in DIA mode exhibits a significant enhancement (>50 fold) in fragment ion signals, providing richer and higher-quality fragment information. Additionally, in Figure 2b, we compare the identified fragment ions of Cyt C from a single tandem mass spectrum acquired using the DIA mode with those from three tandem mass spectra collected in the DDA mode (for precursor ions at charge states of 7+, 8+, and 9+). Furthermore, Figure 2c compares the corresponding sequence coverage maps of Cyt C, in which sequence coverage rate was improved from 7.7% to 24.3% (>3-fold improvement). Both DIA and DDA approaches discovered that Cyt C was acetylated and possessed a heme group. Upon comparing the orange-highlighted position scopes that harbors PTMs, it becomes evident that the amino acid position range containing PTMs is more confined in the DIA mode. In this experiment, deconvoluted tandem mass spectra were processed using ClipsMS^[49]^ to assign internal and terminal fragments. Validity of the results was ascertained through isotope fitting of Cyt C and its fragments, as depicted in Figure 2d. The isotope fitting score of Cyt C is 0.908, and those of b- and y-fragments are 0.958 and 0.938, respectively.

**Figure 2.**
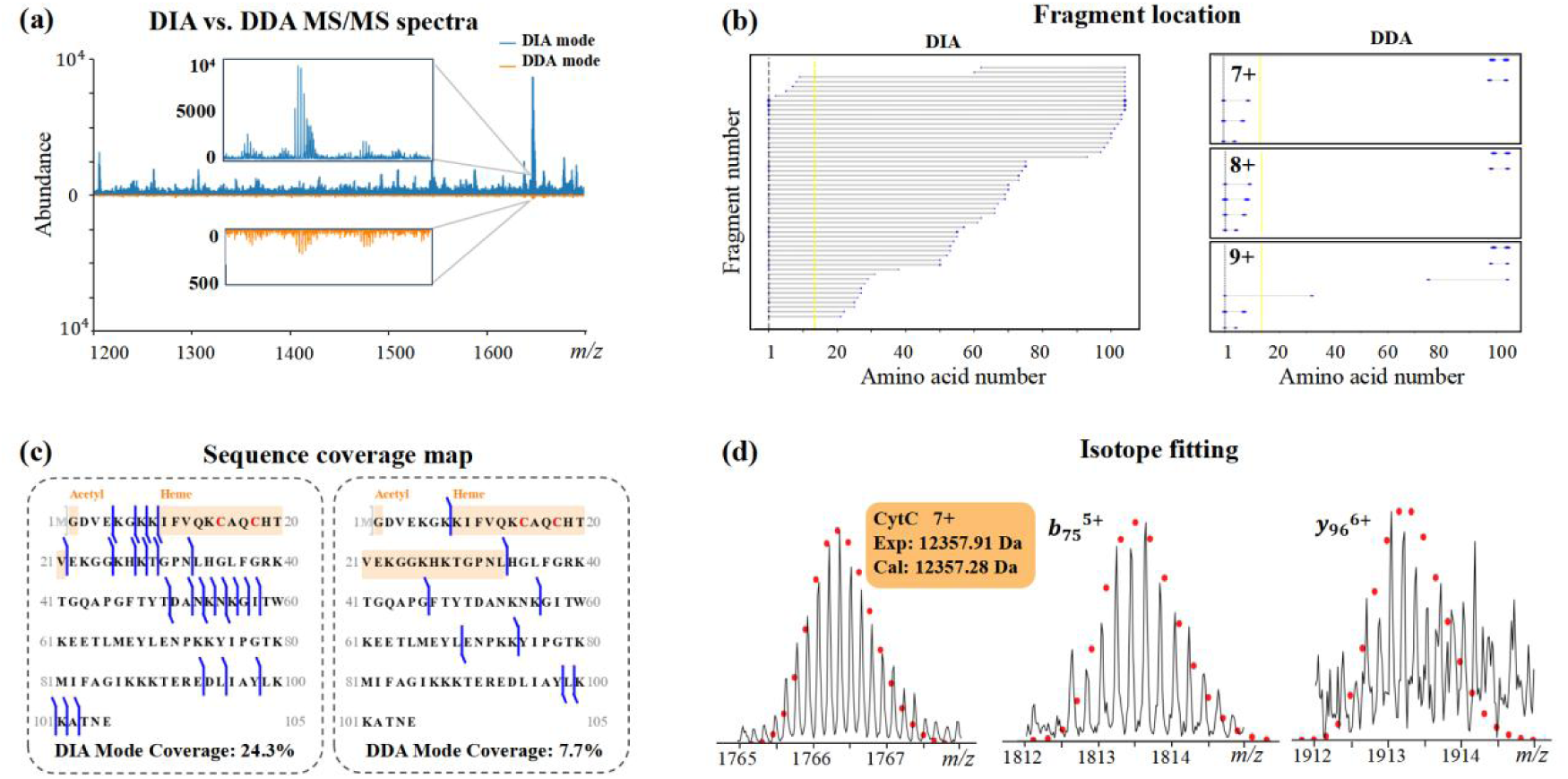
(A) Tandem mass spectra of Cyt C under DIA and DDA modes. (B) Fragment location maps under DIA and DDA modes. (C) Sequence coverage maps using two approaches. (D) Isotope fitting of Cyt C and two of its fragments.

### Analysis of a five-protein mixture

Next, this FW-DIA method was evaluated by analyzing a mixture of five model proteins with molecular masses ranging from 12 to 150 kDa: Cyt C, lysozyme (Lys), beta-lactoglobulin (BLG), serotransferrin (Trf) and immunoglobulin G (IgG). A size exclusion chromatography (SEC) column was used to separate these five proteins under native conditions. Figure 3a plots elution profiles of these five proteins. After deconvoluting tandem mass spectra, elution profiles of fragment ions were acquired using the feature detection module, and these fragment ions were paired with these five protein precursors using the ion paring module. Figure 3b plots a typical tandem mass spectrum (scan #129), in which fragment ions originated from both Lys and Cyt C were presented. Yet, the analysis of their elution profile enables the accurate assignment of precursor ions to their corresponding fragments. As shown in Figure 3b bottom, fragment ions of Cyt C have similar elution profiles (PCC 0.986), and distinct from that of Lys fragment ions (PCC 0.509). The profiles of typical fragment ions were plotted in Figure 3c, which shows that fragment ions would have strong temporal correlations with their corresponding precursors. Effectiveness of the ion paring module was characterized by comparing the protein identification results with or without using this ion paring module. When directly importing DIA data into TDP DDA software without using the ion pairing strategy, the DDA software assumes it is dealing with a single precursor ion and its corresponding fragments. Consequently, all deconvoluted fragment ions were assigned to the precursor ion from the previous MS scan and underwent protein searching. As shown in Figure 3c, ion pairing strategy not only successfully identified all five proteins but also enhanced sequence coverage. It should be noticed that IgG was not treated with reducing agents to break the disulfide bonds, and the light chain and heavy chain were identified in this experiment.

**Figure 3.**
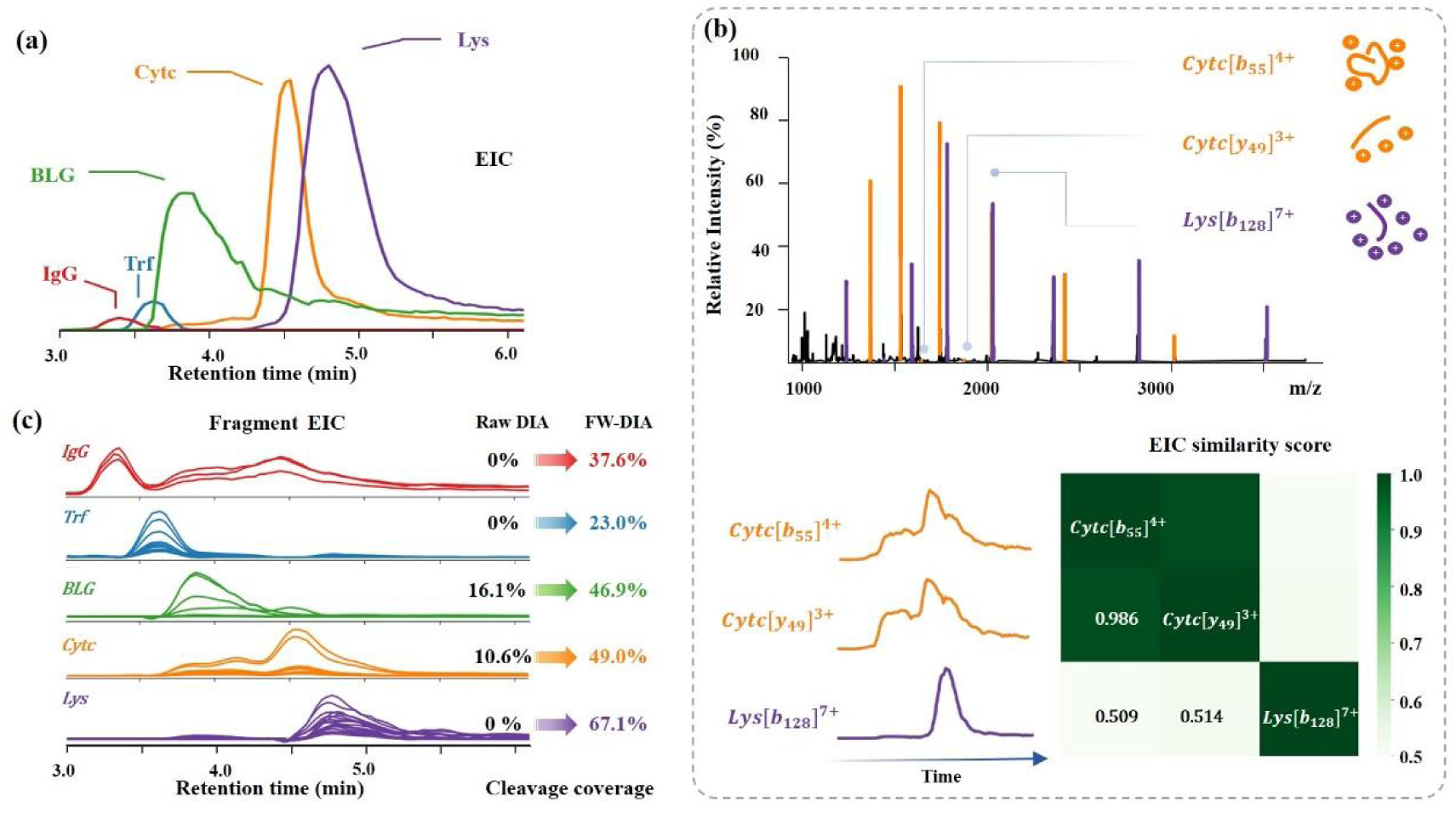
DIA analysis of a five-protein mixture. (a) Elution profiles for five model proteins: Cyt C, Lys, BLG, Trf and IgG. (b) A typical DIA tandem mass spectrum of scan #129, and the corresponding elution profiles plot of fragment ions. The heat-map shows the covariance similarity among the three curves. (c) Elution profiles of fragments, and improved sequence coverage with the FW-DIA approach.

### Analysis of *E. coli* ribosomal proteins

Ribosomes are cellular machinery responsible for protein synthesis. The 50S ribosomal subunit of *E. coli* comprises of 34 ribosomal proteins (RPs), numbered L1 to L36. Notably, L8 is a pentametric stalk complex which composed of L7/L12, where L7 and L12 are nearly identical, differing only in that L7 is acetylated at its N-terminus while L12 is unmodified^[51–53]^. The 30S subunit contains 21 RPs (S1 – S21), and there is no difference between S20 of the small subunit and L26 of the large subunit^[54]^. The FW-DIA method was then applied to the characterization of *E. coli* ribosome proteins. As illustrated in Figure 4a, *E. coli* RPs were first separated by RPLC and subsequently detected by the mass spectrometer. In this investigation, two MS data acquisition methods were used: the conventional DDA mode and the proposed FW-DIA mode. Figure 4b compares the protein identification results using these two methods. A total of 54 RPs (out of 55 RPs) were successfully identified using the FW-DIA method, underlining the elevated sensitivity and robustness of the FW-DIA strategy employed in this study. Figure 4c displays the cleavage coverage rate of each identified RP. Compared with conventional DDA method, this FW-DIA method yielded a higher count of RP identifications, rising from 24 to 54. The maximum sequence coverage achieved was approximately 67%. A recent top-down study^[18]^ of a panel of six standard proteins across different instruments and laboratories revealed an average cleavage coverage for carbonic anhydrase (29 kDa, a protein similar in molecular weight to many ribosome proteins) of approximately 23.5 ± 9.3%. The high sequence coverage achieved in this study was largely facilitated by the full window DIA approach, as well as the machine learning assisted denoising process. Protein sequence coverage rate could be further enhanced using MS instruments equipped with complementary dissociation methods, such as ETD, UVPD and others.

**Figure 4.**
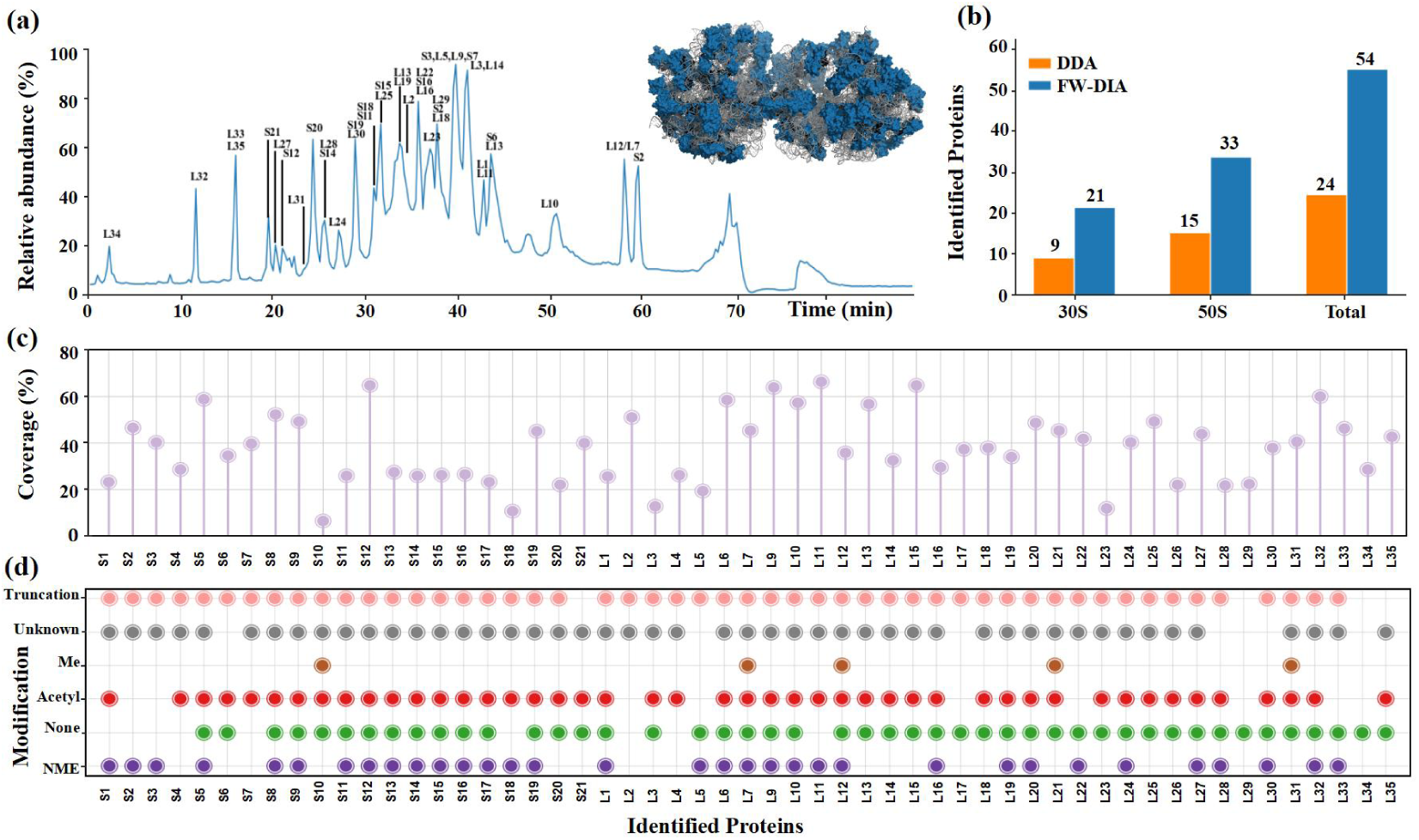
FW-DIA proteomics characterization of *E. coli* ribosomal proteins. (a) RPLC chromatograms of *E. coli* ribosomal proteins. (b) The number of proteins identified using conventional DDA and FW-DIA, respectively. (C) Protein sequence coverage rate of the identified proteins by FW-DIA approach. (D) Identified proteoforms. Me: methylation; Acetyl: acetylation; NONE: no modifications; NME: N-terminal methionine excision.

As shown in Figure 4d, various preteoforms of these RPs were also identified, with N-terminal methionine excision being a common modification among the *E. coli* RPs. The PTMs observed in the RPs of *E. coli*, such as acetylation and methylation, are covalent modifications that can alter structure and function of ribosomes. A number of truncated proteofroms were identified in this study. These truncations may arise from processes such as proteolytic cleavage, RNA splicing, or errors during protein synthesis. The presence of these truncated proteoforms is likely to be biologically significant, as they have the potential to alter protein function, stability, and interaction networks, thereby potentially impacting cellular processes and overall organismal health^[55,56]^ Notably, among these truncated proteins, we found truncations in ribosomal proteins S2, S3, and S4, which are known to form direct contacts with RNA polymerase at its RNA exit site during transcription-translation coupling in *E. coli*. Such truncations could potentially impact this crucial coupling process^[55]^. These identified proteoforms were all confirmed and validated by the isotope pattern fitting module.

Among the 54 proteins identified, the detection of L7, L12, and their associated proteoforms, including methylated variants, was particularly noteworthy. These findings highlight the method’s sensitivity and specificity in detecting both proteins and their PTMs, providing deeper insights into the protein landscape being studied. L7 is identical to L12 except for its acetylated N terminus. Figure 5a shows two groups of peaks, with retention times of ∼58.0 and ∼59.5 minutes, respectively. A closer examination of the mass range region around 1105 Th reveals that each cluster consists of two distinct m/z values, corresponding to their methylated counterparts. According to the elution profile graph, L7, L12 and their proteoforms were detected and clearly resolved using the FW-DIA method. The MS data allowed us to discern the subtle differences between these proteoforms, demonstrating the high-resolution capability of our method in detecting PTMs and protein isoforms.

**Figure 5.**
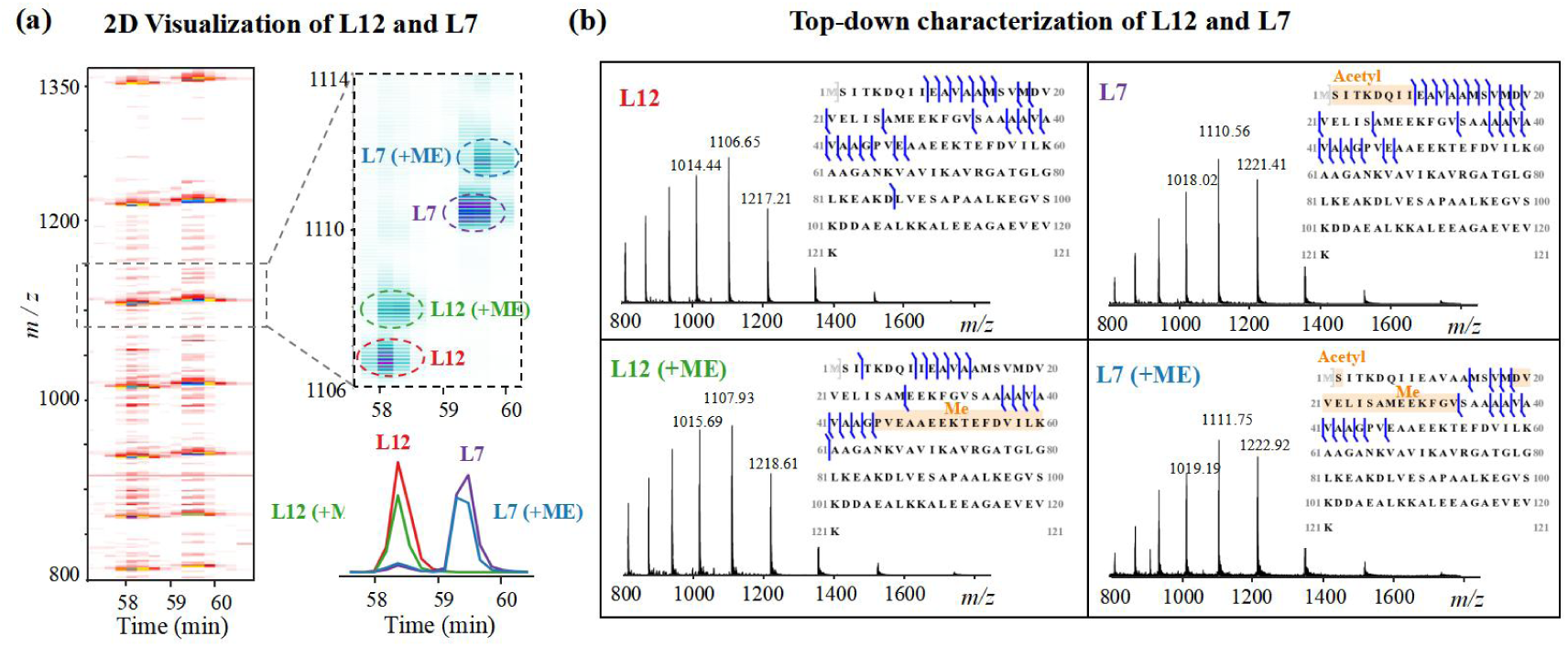
Characterization of L12 and L7. (a) 2D visualization of L12 and L7 in terms of retention time and m/z. Zoomed-in views focusing on the mass range from 1105 to 1115, indicating four different proteoforms. (b) Corresponding MS spectra of L7, L12 and its proteoforms.

## Methods

### Chemicals and materials

Five model proteins: cytochrome C (Cyt C), lysozyme (Lys), beta-lactoglobulin (BLG), serotransferrin (Trf) and immunoglobulin G (IgG), and ribosome proteins were used in this study. Cyt C and BLG were purchased from Macklin Inc. (Shanghai, China). Trf was purchased from SIGMA Inc. (Shanghai, China) and Lys was purchased from Amresco Inc. (Shanghai, China). Trastuzumab (Herceptin) as the model of IgG was purchased from Genentech Inc. (South San Francisco, CA, USA). *E. coli B* strain ribosome was purchased from New England BioLabs Inc. (Ipswich, MA, USA). Deionized water was purchased from Wahaha Co. (Hangzhou, China). All protein samples used in experiments were diluted in 20 mM ammonium acetate at a final concentration of 1 μM.

### Instrument setup

The samples were analyzed using a Xevo G2-XS QTof mass spectrometer (Waters Corporation, Wilmslow, UK) coupled with a Waters Acquity ultra-performance liquid chromatography (UPLC) I-Class system. In the conventional DDA mode, low-energy (6 eV) exact mass precursor ion spectrum was first acquired, which serves as the survey scan. During the DDA MS/MS step, the five most intense precursor ions were isolated (with a window size of 3 Th) and activated using the HCD method. On the other hand, the mass spectrometer alternates between two functions in the FW-DIA mode. The first function (the survey scan) acquires low-energy (6 eV) exact mass precursor ion spectra. In the second function (MS/MS scan), all ions in the mass range would undergo HCD in the collision cell, and elevated-energy exact mass fragment ion spectra were then obtained (Figure 1a).

A ACQUITY UPLC protein BEH SEC column (2.1 mm × 150 mm, 1.7 µm, 200 Å) was used for size exclusion chromatography coupled to the mass spectrometer, when analyzing the five-model protein mixture. Ammonium acetate (AMA) solution was used as the mobile phase with a flow rate of 0.1 mL/min. The electrospray ionization (ESI) voltage was set to 2 kV, while the sampling cone voltage was set to 40 V. The cone gas flow was set to 50 L/h, whereas the desolvation gas flow was 600 L/h. Source temperature and desolvation temperature were set to 150 and 400 °C, respectively. After optimizing the experiment parameters, the MS collision energy was ramped from 70 to 105 eV. Mass spectra were acquired in the mass range of 500-8000 Th.

A UPLC protein BEH C4 column (2.1 mm × 50 mm, 1.7 µm, 300 Å) was used when analyzing *E. coli* ribosome proteins. In the reverse phase liquid chromatography (RPLC) system, phase A (0.1% FA aqueous solution, V/V) and phase B (0.1% FA in ACN, V/V) were used for the experiments. LC-MS/MS runtime was set to 75 min with a flow rate of 0.2 mL/min. Mobile phases A and B were used for gradient elution: 5% B for 5 min, 5% - 42% B for 60 min, 42% - 95% B for 5 min, keeping 95% B for 5 min. Mass range of the mass spectrometer was set at 500 - 3000 Th, and the HCD collision energy was ramped from 20 to 70 eV after optimization. Both DDA and DIA experiments were carried out for comparison. In the DDA experiment, 0.2 and 0.1 s were found to be the optimized scan times for the survey and MS/MS scans, respectively. In the DIA experiment, the survey scan time was set at 5 s, and the MS/MS scan time was set at 1 s after optimization.

### DDA data analysis

After collecting DDA data from the Waters Xevo G2-XS QTof, the raw files (.raw) were converted into HUPO-PSI compliant mzML files using MSConvert^[57]^ with the “peak picking” filter. Subsequently, the mzML files were subjected to spectral deconvolution via TopFD^[47]^ The resulting msAlign files were then subjected to a top-down search engine software: TopPIC (version 1.6.2) with the objective of identifying and characterizing proteoforms from a top-down perspective. The *E. coli* DDA data were searched against the *E. coli* strain B database downloaded from Uniprot. No variable modifications were specified, and the error tolerance for precursor and fragment masses was set to 15 ppm. Results were filtered using a 1% PrSM-level E-value, and 1% proteoform-level E-value.

### Training data collection and data preprocessing model construction

The CNN model was trained using ESI mass spectra data collected on the QTof mass spectrometer. The raw data underwent a two-stage preprocessing approach. First, spectral deconvolution was performed using FLASHDeconv. Following deconvolution, a manual review was conducted to identify which mass range intervals contained signals and which did not. Subsequently, each spectrum was divided into small windows of length 100. If a window contained a protein peak, it was labeled as a protein signal; otherwise, it was labeled as noise. Due to the abundance of noise in the data, the noise samples were manually selected to balance the number of noise and protein signals in the training set. This manual review ensured high-quality training data, evenly covering the entire mass range from 0 to 7000. Finally, the training dataset comprised 23,542 MS1 data samples and 9,700 MS2 data samples. The MS2 dataset was augmented from the MS1 dataset to account for differences in noise and signal patterns between the two data types.

Two separate CNN models were developed: one for MS1 data and another for MS2 data. This separation was necessary because the characteristics of noise and protein signals differ between MS1 and MS2 spectra. CNNs were chosen to identify complex patterns that distinguish protein signals from noise in MS data. Each CNN model processes training data by sequentially extracting features. Starting with three convolutional layers that use filters to identify local patterns, the models progressively learn more complex features. Max pooling layers reduce dimensionality, and a flattening layer prepares the data for fully connected layers, which combine and refine these learned features. L2 regularization and dropout layers prevent overfitting, and a final sigmoid output layer predicts the probability of an input spectrum being a protein peak or noise.

To minimize false negatives while optimizing both the true positive rate (TPR) and true negative rate (TNR), classification thresholds were carefully determined for both the MS1 and MS2 models. This approach effectively balances sensitivity and specificity, enhancing the models’ accuracy in distinguishing protein signals from noise.

### DIA data analysis

Similar to the DDA data, the raw DIA data were first converted into mzML files in profile format using MSConvert, including both MS and MS/MS data. All resulting mzML files were processed through a de-noising model. After denoising, MSConvert was used again to convert the denoised mzML files into centroided format using the peak-picking option. Three deconvolution software tools then detected features, and the results were merged based on molecular weight, continuous m/z values, and retention-time spans. In DIA mode, since multiple precursor ions were fragmented simultaneously, it is essential to match each fragment ions with its corresponding precursor ions^[58]^. This procedure calculates the similarity between the elution profiles of precursor and fragment ions, producing msalign files for subsequent identification. Protein identification was performed using TopPIC as the search engine, and results were filtered based on theoretical isotope fitting and mass alignment. Identified proteoforms were retained if they had a quantified number of matched fragments with an isotope fitting score of at least 0.8.

## Discussion

Due to the presence of multiple charge states for protein ions in ESI mass spectra, a novel DIA approach was developed for TDP without employing isolation windows for precursor ion selection. Instead, all ions eluted from the LC system were subjected to fragmentation, resulting in significantly enhanced fragment ion intensity and increased fragment diversity. In the FW-DIA process, mass spectrum deconvolution was enhanced by a machine learning based denoising module, as well as integrating the outcomes of three complementary deconvolution methodologies. This approach generates a dependable deconvoluted mass list and feature data, reducing the likelihood of mass artifacts. Furthermore, an ion pairing strategy was developed to match precursor with fragment ions. The unbiased nature of DIA allows for the detection of a wider range of fragment ions, reducing the precursor selection bias. As a result, FW-DIA exhibited increased number of identified proteins compared to DDA, while also providing improved sequence coverage and more accurate identification of PTM sites. Performances of FW-DIA were demonstrated using a five-protein mixture in native conditions (SEC plus native ESI) and *E. coli* ribosomal proteins in high chromatographic resolution mode (RPLC plus conventional ESI). This FW-DIA method not only provides boosted proteoform identification performances, but also enables the application of DIA based TDP to most commercial MS instruments. Besides LC-MS based platform, this FW-DIA is also compatible with the fast-developing separation techniques, such as ion mobility and capillary electrophoresis. As these methods for TDP also continues to evolve, we anticipate that DIA will play an increasingly important role in the analysis of complex biological samples, enabling deeper insights into proteoform biology and function.

## Data availability

All MS proteomics data have been deposited to the ProteomeXchange Consortium (http://proteomecentral.proteomexchange.org) via the iProX repository with the project ID PXD057331.

## Code availability

The source code and the pretrained model for FW-DIA are available in the Github repository (https://github.com/grabriellechenchenchen/FW-DIA) for academic use. The code on Github serves as an esay-to-use tool for running the data processing procedure for FW-DIA.

## Acknowledgements

This work was supported by NNSFC (22374009). We also thank the Analysis & Testing Center of Beijing Institute of Technology for MS instrument support.

## Contributions

W.X. contributed to conceptualization, supervision, management, manuscript reviewing and editing. C.S. developed the data processing procedure, analyzed the data, and drafted the manuscript. W.Z. contribute to the MS experiments, and revised the manuscript. M.Z. and M.H. contributed to manuscript reviewing and editing.

## Ethics declarations

