## Supplementary Data for "A Full Window Data Independent Acquisition Method for Deeper Top-down Proteomics"

***Supporting information***

Chen Sun[1], Wenjing Zhang[1], Mowei Zhou[2], Martin Myu[3], Wei Xu[1*]

[1]School of Medical Technology, Beijing Institute of Technology, Beijing 100081, China

[2]Department of Chemistry, Zhejiang University, Hangzhou 310058, China

[3]Institute of Food Safety, Chinese Academy of Inspection and Quarantine, Beijing 100176, China

*Corresponding Author:
Wei Xu
School of Medical Technology
Beijing Institute of Technology
Haidian, Beijing, 100081, China


### Top-down DIA data preprocessing model

Top-down mass spectrometry is a powerful approach for analyzing intact proteins, but the resulting data are characterized by high mass, multiple charges, and broad peak widths, making the spectra increasingly complex. Furthermore, the DIA mode of TD-MS exacerbates peak overlap, rendering the spectra even more challenging to analyze. Additionally, top-down deconvolution software is heavily influenced by data quality, leading to high false positive rates in deconvolution results. To address these challenges, convolutional neural network (CNN)-based models were developed to differentiate between noise and authentic protein signals in TD-DIA MS data (Supplementary Fig. 1). Utilization of this model enables the efficacious elimination of noise interference peaks from spectral data, thereby enhancing the preprocessing of TD proteomics data and culminating in augmented efficiency and confidence in subsequent qualitative analyses.

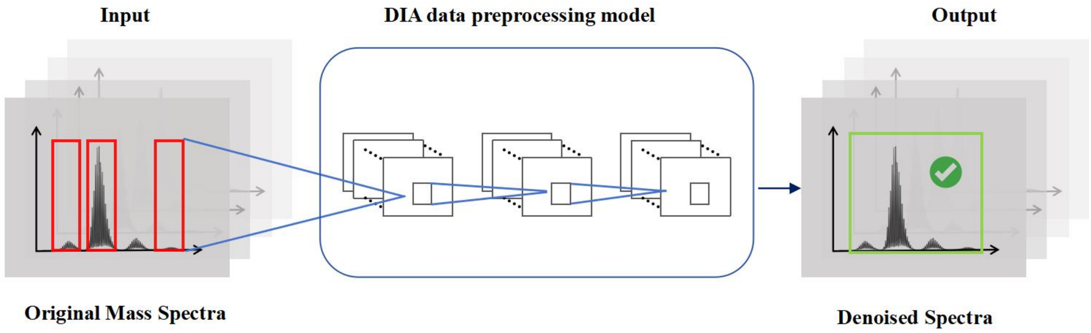


**Supplementary Fig. 1**Illustration of the DIA data prepossessing model.

The proposed framework consists of two distinct models, tailored to handle MS1 and MS2 data separately. The reason for training two models separately is that there are differences in the noise characteristics and true protein signal distributions between MS1 and MS2 data, necessitating the development of specialized models for each stage. The model for MS1 data preprocessing procedure was trained on a manually labeled dataset comprising 23,542 MS1 samples, achieving a validation accuracy of 0.957 and a loss of 0.191. Given the critical objective of minimizing false negatives (i.e., ensuring that true protein signals are not missed), while simultaneously optimizing the true positive rate (TPR) and true negative rate (TNR), a classification threshold for the MS1 model was determined to be 0.3 (Supplementary Fig. 2). This threshold strikes an optimal trade-off between TPR and TNR, effectively balancing sensitivity and specificity in the model.

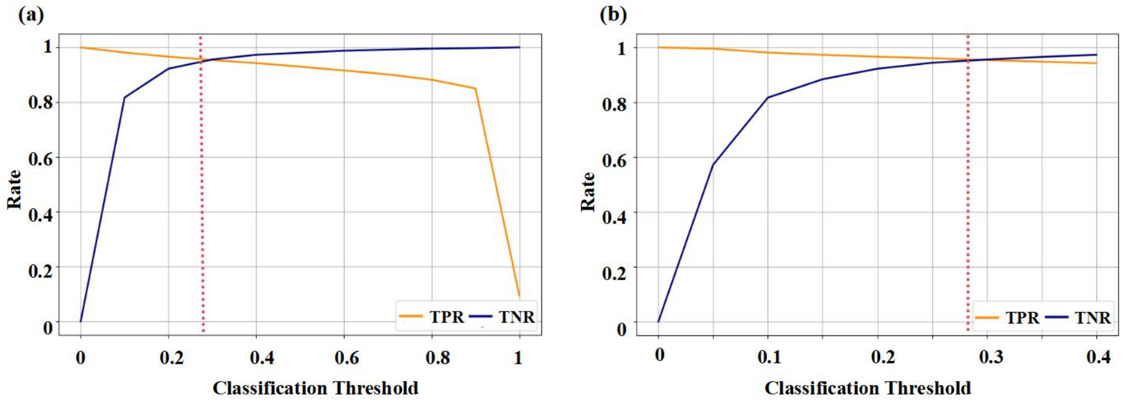


**Supplementary Fig. 2** (a) The trends of TPR and TNR when the classification threshold changes from 0 to 1; (b) The trends of TPR and TNR when the classification threshold changes from 0 to 0.4.

The dataset employed for training the MS2 level data preprocessing model was constructed by augmenting the MS1 dataset with an additional 9,700 MS2 samples, yielding a final training set comprising approximately 35,000 samples. This augmentation was necessitated by the distinct noise and signal patterns present in MS2 data, which, although sharing similarities with MS1 data, also exhibit notable differences. The MS2 model was also trained and evaluated, focusing on maintaining high TPR and TNR, while maximizing the identification of noise. The model achieved a test accuracy of 0.9009 and a validation accuracy of 0.827, with a loss of 0.24976. To reduce the false positive rate (FPR), the classification threshold of the MS2 model was 0.1, which intentionally prioritized a higher specificity to ensure the detection of true protein signals.

As shown in Supplementary Fig. 3, the data preprocessing models can effectively distinguish noise from protein/peptide signals. Comparing the original and denoised peak intensity distribution histograms, it is found that the low-intensity noise peaks are significantly reduced, while the high-intensity true signal peaks are basically preserved. This indicates a successful removal of noise, enhancing the signal-to-noise ratio and improving data quality. After applying the MS1 and MS2 models as an initial data preprocessing step, the false positive rate in the subsequent deconvolution step was significantly reduced. Supplementary Fig. 4 illustrate the number of masses and the corresponding mass distribution detected in a five-protein mixture sample and in the *E. coli* ribosomal protein sample using FlashDeconv under various noise removal conditions. Compared to traditional methods that employs a threshold to filter out noise, the use of deep learning models enables the deconvolution of a broader and more enriched range of masses, while threshold-based approaches (i.e. BPI > 0.005) may result in the loss of low-abundance signals. This improvement underscores the efficacy of the deep learning models in effectively filtering out noise, thereby enhancing the overall performance of the proteomic analysis pipeline. The integration of these models into the preprocessing workflow offers a robust solution to the challenges posed by noise in DIA proteomics data, facilitating more accurate and reliable protein identification.

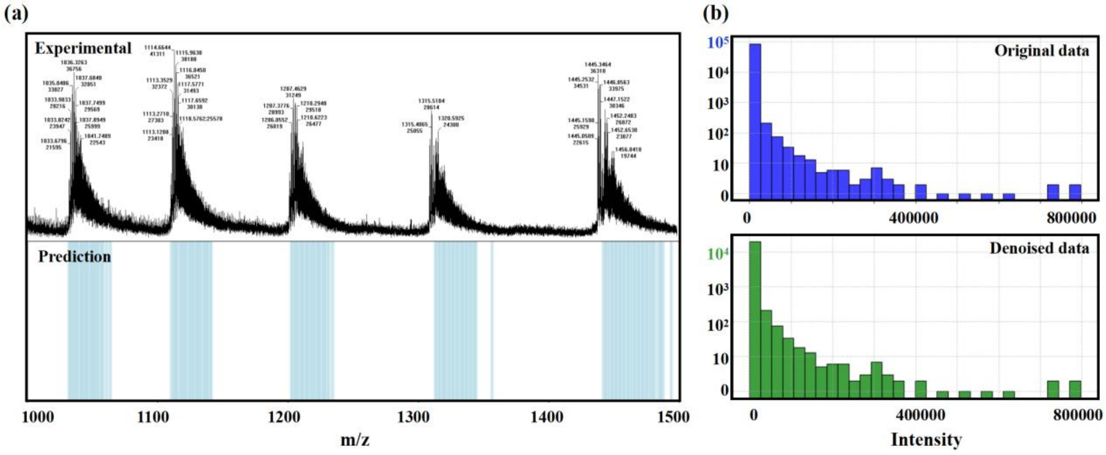


**Supplementary Fig. 3** (a) Experimental mass spectrum and visualization of the model's prediction results. (b) Original and denoised peak intensity distribution histograms.

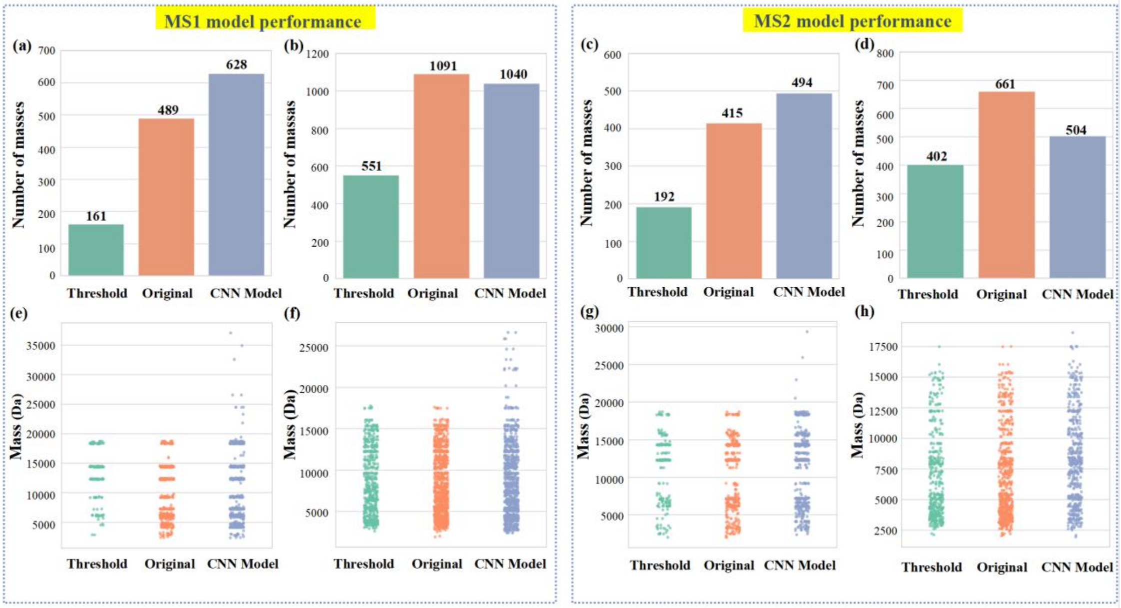


**Supplementary Fig. 4** (a) The number of masses in a five-protein mixture MS1 sample using MS1 preprocessing model detected by FlashDeconv under different noise removal conditions: threshold (only peaks with intensities above 0.005 times the base peak), original data without noise removal, and noise removal by CNN models; (b) The number of masses in an E. coli ribosome protein MS1 sample using MS1 preprocessing model detected by FlashDeconv under different noise removal conditions; (c) The number of masses in a five-protein mixture MS2 sample using MS2 preprocessing model detected by FlashDeconv under different noise removal conditions; (d) The number of masses in an E. coli ribosome protein MS2 sample using MS2 preprocessing model detected by FlashDeconv under different noise removal conditions; (e) Mass distribution in a five-protein mixture MS1 sample under various noise removal processing methods using MS1 preprocessing model; (f) Mass distribution in an E. coli ribosome protein MS1 sample under various noise removal processing methods using MS1 preprocessing model. (g) Mass distribution in a five-protein mixture MS2 sample under various noise removal processing methods using MS2 preprocessing model; (h) Mass distribution in an E. coli ribosome protein MS2 sample under various noise removal processing methods using MS2 preprocessing model.

**Result verification by Isotopic pattern matching**

After receiving the TopPIC results, an isotopic pattern fitting technique was used to verify the detected protein fragments and evaluate data quality. As illustrated in Supplementary Fig. 5, ribosomal protein S15 was identified with a high confidence p-value. ${y_{57}}^{8+}$ and ${y_{61}}^{9+}$ of S15 are identified, the similarity score between the theoretical isotopic pattern and the actual peaks with Pearson correlation coefficients are 91.047 and 85.336 respectively, indicating a highly degree of concordance. The m/z values of all peaks, however, show a slight fluctuation of less than 0.1 Th, which is larger than the theoretical value, as shown in Supplementary Fig. 5 C and F. This could be because to the instrument's mass axis was slightly misaligned and requires additional instrument calibration.

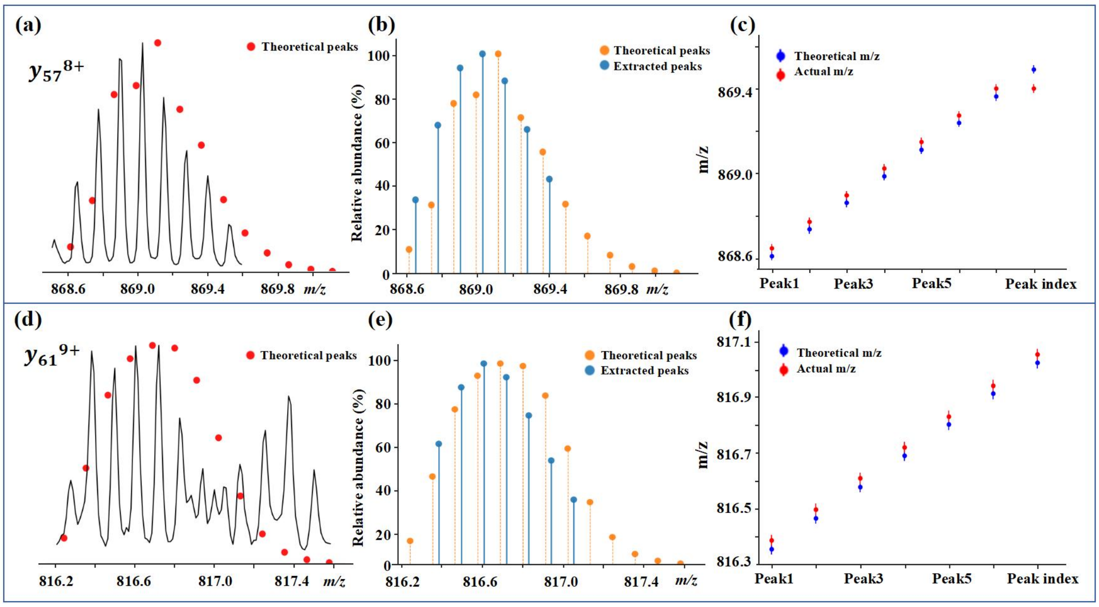


**Supplementary Fig. 5** (a) and (d): two Y ion fragments of S15 and their theoretical isotope simulation, respectively; (b) and (e): the intensity comparison between the actual and theoretical centroided peaks; (c) and (f): peak position comparison.

**DIA experiment parameter optimization.**

Instrument parameters for top-down MS studies were carried out using a Waters G2-XS Tof system. The $MS^E$ acquisition mode was selected, and the parameters related to the MS/MS acquisition, such as collision energy ramp, were optimized for better identification. Here, we optimized the experiment settings for cytochrome C. As shown in Supplementary Fig. 6 a, three consistent collision energies were first set: 50 eV, 70 eV, and 100 eV, and a ramping energy was set from 60 eV to 90 eV. Evaluation of the experiment's results was done by observing the number and the mass distribution of fragments. As depicted in Supplementary Fig. 6 b, the fragments obtained under condition 3 setting (100 eV) were significantly fewer than those obtained under other conditions, which was contrary to expectations. Higher energy does not necessarily result in an increased number of fragments. Next, although the condition 1 setting (50 eV) earned the most fragments among three consistent collision energies, the abundance of the fragments is much lower than the 70 eV condition. In addition, the fragmentation efficiency under the condition of collision energy ramping from 60 to 90 eV was not as good as that of the 70 eV. As a result, 70 eV was chosen to be the optimal collision energy for the experiment of cytochrome C.

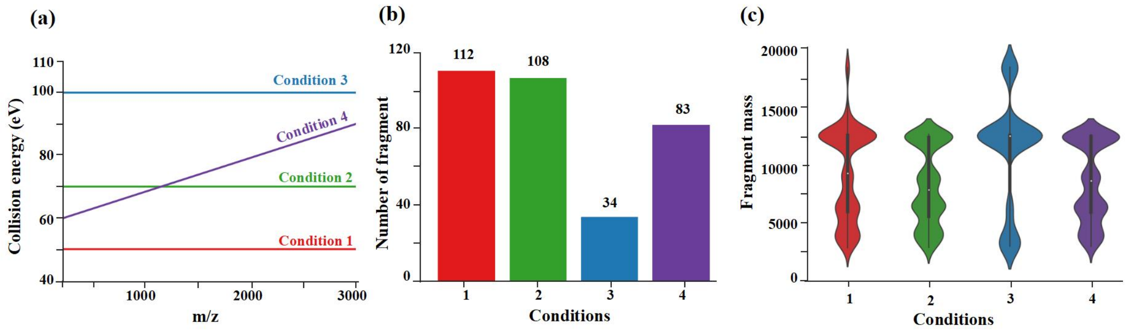


**Supplementary Fig. 6** (a) Different collision energy settings; (b) Collision energy vs the number of deconvoluted fragments; (c) Fragments masses distribution among different collision energy conditions.
